# Subadult macaws (*Ara glaucogularis*) copy better than adults but are less likely to imitate in problem-solving tasks

**DOI:** 10.1101/2025.09.20.677516

**Authors:** Esha Haldar, Deepshika Arun, Auguste M.P. von Bayern

**Affiliations:** Max-Planck-Institute for Biological Intelligence, Munich; Ludwig-Maximilians-Universität, Munich; Comparative Cognition Research Center, Loro Parque Fundacion, Tenerife, Spain; University of Zurich, Zurich, Switzerland

**Keywords:** motor imitation, copy, parrots, social learning, age effect, two-action task, bidirectionally controlled tasks

## Abstract

Copying via imitation (replicating others’ bodily actions) or emulation (replicating others’ action goals) underpins the cultural transmission of social norms and technical skills in humans. In this study, we investigated whether blue-throated macaws (*Ara glaucogularis*) are capable of copying conspecifics using two problem-solving devices: ‘door’ and ‘plug’. The ‘door’ device could be solved by sliding a panel either *UP* or *DOWN*, while the ‘plug’ device required removing a stopper via either a *PUSH* or *PULL* action. A control group (N=5), which received no demonstrations, consistently preferred a single solution for each device— *DOWN* for ‘door’ and *PULL* for ‘plug’. In the subsequent test, adult (N=5) and subadult (N=5) macaws from the test group observed demonstrators (N=2) performing the *non-preferred* action for each device. Significantly more test birds *copied* the demonstrated action (*UP)* for ‘door’, but not *PUSH* for ‘plug’, because the subadults were less likely to *imitate* the *PUSH* action than the adults. However, the subadults took significantly less time to approach and manipulate the devices than the adults. Overall, our study shows that macaws can copy conspecifics in problem-solving tasks, which facilitates information transfer and transmission of foraging techniques, a prerequisite for foraging culture to arise.

## INTRODUCTION

Social learning (learning by observation of others) supports the emergence of locally adaptive behavioural patterns (Galef, 1995) and therefore also cultural transmission of behaviour within different animal populations (van Schaik et al., 2003; Whiten, 2021). By exploiting the knowledge of experienced individuals, animals can save time and energy otherwise required for trial-and-error learning, with potentially significant fitness benefits (Slagsvold & Wiebe, 2011). Consequently, scientists have long sought to understand which aspects of conspecific behaviour and environmental cues animals attend to, so as to learn effectively from each other through observation (C. M. Heyes, 1994; Zentall, 2006). Several mechanisms that may underpin social learning and vary in complexity have been proposed. In *social facilitation,* for example, the observer gets motivated to approach and solve a problem from observing a conspecific (Zajonc, 1965), in *stimulus* and *local enhancement*, the observer learns ‘what’ (i.e. which object) and ‘where’ (i.e., which location) in the environment to focus on (Zentall, 2006), while in *affordance learning*, the observer learns about an object’s properties and functions through the actions of a demonstrator (Whiten & Ham, 1992). When *copying*, the subject learns to copy a) the *means* of an action (*imitation*), b) the end state, i.e., goal of the action (*emulation*) or c) the goal of the object movement (*object movement reenactment*) (Whiten & Ham, 1992).

*Imitation*, where an observer “learns and reproduces ‘something’ about the topography of the model’s body movements” (C. Heyes, 2023) is considered the most sophisticated social learning process. By contrast, in *emulation* or *object movement reenactment*, the observer learns about the events occurring in the environment as a result of the action (Whiten and Ham 1992). While imitation is necessary for faithful copying of opaque cultural behaviours like goalless actions or body movements (e.g., gestures or forms of dance) in humans (reviews: Csibra & Gergely, 2006; Gergely & Csibra, 2020)), emulation is thought to have underpinned learning novel instrumental tasks (e.g., tool use) during human cumulative cultural evolution (Gergely and Csibra 2006). It also is considered the main mechanism of cultural learning in non-human primates (Tomasello et al., 1994). Yet, to distinguish between these forms of copying has often proven to be difficult, as the behaviour of the observer and the model are very similar in each case. Both copying mechanisms can be distinguished from other forms of social learning (like *stimulus or local enhancement*) by using the two-action task (Voelkl & Huber, 2000), and its variant, the ‘bidirectional control’ method (Fawcett et al., 2002). Both methods have become the benchmark paradigms for studying imitative learning (Zentall, 2006). In the ‘two-action task’, subjects observe a trained conspecific extract a reward by either performing one of two possible actions (e.g., push vs. pull a stopper) or one action in two particular ways (e.g. remove a lid by hand vs. mouth). In contrast, in the ‘bidirectional’ approach, the problem is solved by executing the task in one of two opposing directions (e.g., slide a panel left vs. right, or push a panel up vs. down). If the subjects perform the action using the observed method or in the same direction as the model, they are considered to have successfully copied their conspecifics. Alternative social learning processes, such as stimulus or local enhancement, are controlled for in these tasks because the same object (e.g. a panel or plug) can be manipulated in two equally possible ways; hence mere attraction to the location or the stimulus does not ensure a match between the observed and executed actions. Two action studies in marmosets (Callithrix sp.) (Voelkl & Huber, 2000), budgerigars (*Melopsittacus undulatus*) (C. Heyes & Saggerson, 2002), pigeons (*Columba livia*) (Zentall et al., 1996), Japanese quail (*Coturnix japonica*) (Akins et al., 2002), common ravens (*Corvus corax)* (Loretto et al., 2020), showed that *imitation* is the underlying mechanism for such animals to socially learn the actions from their conspecific. (Watson et al., 2018). However, unlike two-action tasks, *imitative* and *emulative* social learning mechanisms cannot be fully teased apart using only bidirectional control tasks, as the subject can learn about the end-state of manipulation from the final object position rather than the action itself, risking the misinterpretation of *object movement reenactment* (an *emulative* process) as *imitation*. Hence, in absence of additional ghost-control groups, where the subjects observe the direction of manipulation being demonstrated without the conspecific manipulating the object (e.g., via an invisible mechanism like using a fishing line or a magnet (Fawcett et al., 2002; Miller et al., 2009; Mioduszewska et al., 2020)), a bi-directional task can only provide evidence for *copying* without distinguishing between *imitation* and *emulation* skills in the tested animals.

Parrots are large-brained birds widely renowned for their vocal imitation ability (Benedict et al., 2022; Chakraborty et al., 2015; Krasheninnikova et al., 2024). They exhibit an extended juvenile period (Salinas-Melgoza & Renton, 2007; Smeele et al., 2022) throughout which the young birds remain associated with their parents providing them with ample opportunity to learn from them. Parrots also typically forage in large fission-fusion flocks (Bradbury & Balsby, 2016) and likely gather information (e.g. on novel food sources or on foraging techniques) from conspecifics etc. Thus, parrots are interesting candidates for investigating social learning, particularly in young individuals, their preferred method of gaining social information (whether high fidelity *copying* vs. low fidelity *enhancement* techniques) remains unclear and hardly studied. In the only two ‘two-action’ studies conducted on parrots, budgerigars (*Melopsittacus undulatus*) belonging to the *Psittaculidae* family showed consistent (C. Heyes & Saggerson, 2002) *copying*, while kea (*Nestor notabilis*), a *Strigopidae* family member, demonstrated no significant tendency to copy trained conspecifics (Suwandschieff et al., 2023). This indicates that social learning strategies may vary across parrot species or families. The goal of the present study was to investigate the copying ability of a further parrot species, blue-throated macaws (*Ara glaucogularis*), a New World parrot species belonging to yet another parrot family, i.e., *Psittacidae*). We investigated whether they could copy a trained conspecific in a bidirectional and a two-action task. Blue-throated macaws are a critically endangered species (Hesse & Duffield, 2000) whose numbers are limited even in captivity, so it was not possible to test an additional ghost control group. Hence, we addressed their *copying* skills without distinguishing between *imitation* from *emulation* for the bi-directional task. Previous studies have shown that captive-bred blue-throated macaws were capable of imitating intransitive, goal-less actions displayed by a conspecific (Haldar et al., 2024, 2025), which is considered as an advanced socio-cognitive skill (Custance et al., 1995) and potentially plays important social functions like enhancing group cohesion and theoretically enabling cultural transmission of gestures (Haldar et al., 2024). However, their social learning ability has not yet been tested in the context of physical problem-solving tasks involving *transitive* actions, i.e. actions involving *goals* or *objects* (Chiavarino et al., 2013). Social learning of transitive actions is more relevant for adapting to a species’ physical rather than a social environment (Caldwell & Whiten, 2002) and this is what the present study aims to focus on.

We used two experimental devices, one consisting of a bidirectionally controlled task and the other of a two-action task, to test the macaws’ social learning abilities in problem-solving, involving actions directed at physical objects. Slightly deviating from the classical two-action/bi-directional task protocol because of our limited sample size, we did not test two ‘social’ groups observing two ways of demonstration. Instead, we first collected baseline data for the natural tendency of macaws from a control (‘asocial’) group. N=5 control subjects were tested in the two problem-solving tasks without observing any demonstrations, to check whether the birds can solve the problem individually by trial and error using any preferred action. The first task, a ‘door’ device (Watson et al., 2018), involved pushing a sliding shutter *UP or DOWN* to open a reward slot providing access to a box containing walnut pieces. The second task, a ‘plug’ device, involved *PUSH*ing in or *PULL*ing out a circular cork (C. Heyes & Saggerson, 2002), opening a similar slot enabling access to food in a box. After establishing the natural preferred responses of the control group in the two devices, we tested the remaining subjects (N=10) for the non-preferred actions, demonstrated by conspecific models in a social condition, If an animal has a naturally preferred action among the alternatives, testing for the non-preferred action (highly improbable to occur) can provide irrevocable evidence for social learning against trial and error learning (Voelkl & Huber, 2000).

Social learning is known to be influenced by a variety of environmental and social factors (see (Lambert et al., 2019) for meta-analysis) as well as life-history traits including phylogeny, ontogeny, dominance rank, age, and sex of the individuals (see (Penndorf & Aplin, 2020) for meta-analysis). Concerning age, social learning in foraging contexts does not seem to differ between juveniles and adults across non-human animals (Penndorf & Aplin, 2020) but juvenile animals tend to be more exploratory and less neophobic, as well as picking up new behaviours faster, compared to adults (Tennie et al., 2010; Zohar & Terkel, 1995) as seen in many birds (Greenberg & Mettke-Hofmann, 2001) including parrots (Auersperg et al., 2015; Greenberg & Mettke-Hofmann, 2001). We examined whether age affected social learning of problem-solving tasks in blue-throated macaws. Thus, we divided the test individuals into two age groups, i.e., sub-adults and adults, and hypothesised that young individuals would show more neophilic tendencies than older adults in these tasks.

Finally, we wanted to pilot whether blue-throated macaws would copy irrelevant tasks that have no function in solving a given problem-solving task. Studies have shown that human children imitate irrelevant actions demonstrated by adults even when an action offers no discernible evidence of serving a function in achieving a given task (Horner & Whiten, 2005), out of cultural-normative habit or to gain social affiliation - a behaviour termed ‘overimitation’ by Lyons and colleagues (Lyons et al., 2007). Once thought uniquely human, rudimentary forms of overimitation have been observed in companion dogs, imitating irrelevant tasks when demonstrated by care givers, a tendency likely shaped by domestication and enculturation (Huber et al., 2018; Mackie et al., 2025). In our study, we introduced two causally irrelevant tasks which were performed before the relevant tasks in a sequence. For the ‘door’ device, the irrelevant action was to rotate once (i.e., a conspicuous behaviour) on the perch before manipulating the door and for the ‘plug’ device, the demonstrator pushed a buzzer with his foot, adjacent to the perch, before manipulating the knob of the plug. We hypothesised that if the subjects imitated the causally irrelevant task, this would indicate that macaws ‘*overimitate’* their conspecifics, which may serve functions in social bonding similar to humans (Lyons et al. 2007). Overall, our study aimed to investigate copying skills of blue-throated macaws in problem solving tasks using two-action procedure and evaluate age effects on social learning in this species.

## MATERIALS AND METHODS

### Subjects (N=10 test; N=5 control)

Blue-throated macaws (*Ara glaucogularis*) (N=15), ranging in age from 0.5 to 15 years (**see Table 1** for individual details) housed at the Max-Planck Comparative Cognition Research Station integrated into Animal Embassy/Loro Parque zoo, Tenerife took part in this study. All subjects of this critically endangered parrot species according to IUCN red list of species were bred by Loro Parque Fundación and were hand-raised before being socialized in groups of conspecifics. They were well habituated to interacting with human experimenters and participating in cognitive studies (see Supplementary Information). N=5 subjects from 0.5-4 years were assigned to the subadult age group, whereas N=5 subjects above 5 years were considered as adults. We considered the age of each subject at the beginning of their testing period and kept it constant for the analysis as testing did not exceed three months with any subject. Prior to the experiment, all subjects had received basic training to enter the test compartment and sit on a perch via positive reinforcement (clicker and food). N=5 randomly selected individuals served as the control group for establishing the baseline preference, while the remaining N=10 birds were assigned as test subjects (see **Table 1)**. Two of the five control subjects served as demonstrators after they had terminated their control sessions and demonstrated their preference of actions/directions. The 10 test birds were pseudo-randomly assigned to one of the two models, counterbalancing the number of subadult and adult individuals to test for the non-preferred actions on the two devices.

**Table 1.** Details of subjects used in the study and their responses in the two tasks.

| Species | Individual | Age (years)/ Sex (Male=M or Female=F) | 'Door' response (Demonstrated action marked by 'D') | 'Plug' response (Demonstrated action marked by 'D') | Group | Demonstrator |
| --- | --- | --- | --- | --- | --- | --- |
| *Ara glaucogularis* | Wanda | 15/ F | UP (D) | Mixed | Adult Test | Lady |
| *Ara glaucogularis* | Charlie | 8/ M | UP (D) | Mixed | Adult Test | Lady |
| *Ara glaucogularis* | Marvel | 3/ F | DOWN | PULL | Subadult Test | Lady |
| *Ara glaucogularis* | Iron-man | 1/ M | UP (D) | PULL | Subadult Test | Lady |
| *Ara glaucogularis* | Pickle | 0.5/ M | UP (D) | PUSH (D) | Subadult Test | Lady |
| *Ara glaucogularis* | Huang | 9/ M | UP (D) | PUSH (D) | Adult Test | Sherlock |
| *Ara glaucogularis* | Thor | 13/ M | DOWN | PULL | Adult Test | Sherlock |
| *Ara glaucogularis* | Carrot | 4/ F | UP (D) | PULL | Subadult Test | Sherlock |
| *Ara glaucogularis* | Natasha | 4/ F | UP (D) | PULL | Subadult Test | Sherlock |
| *Ara glaucogularis* | Gargamel | 10/ M | UP (D) | PUSH (D) | Adult Test | Sherlock |
| *Ara glaucogularis* | Pepper | 1/ F | DOWN | PULL | Subadult Control | - |
| *Ara glaucogularis* | Lady | 8/ F | DOWN | PULL | Adult Control & model | - |
| *Ara glaucogularis* | Sherlock | 15/ M | DOWN | PULL | Adult Control & model | - |
| *Ara glaucogularis* | Long John | 9/ M | DOWN | PULL | Adult Control | - |
| *Ara glaucogularis* | Morty | 4/ M | DOWN | PULL | Subadult Control | - |

### Housing conditions

All parrots were group-housed in semi-outdoor aviaries (see Supplementary Information for details). The birds had 24-hour access to the outside aviary, allowing them to follow a natural light schedule. The subjects were kept in 3 adjacent but separated aviaries which maintained a group of 5-6 individuals each. All birds had ad libitum access to water and mineral blocks and received a ration of fresh fruit and vegetables twice a day along with a seed mix (Loro Parque Ara mix) only in the afternoon. To transport the subjects to the testing rooms, the aviaries were connected with mobile (1mx1mx1m) feeding cages which could be wheeled with ease with the birds inside. The birds were completely used to this procedure and entered the cages to be transported readily and voluntarily.

### Experimental setup

Testing took place in an indoor testing chamber equipped with lamps covering the birds’ full range of visible light (Arcadia 39 W Freshwater Pro and Arcadia 39 W D3 Reptile lamp). Measurements of the testing chamber were 2.5 m × 1.5 m × 1.5 m (height × width × length). A table of measurement 1.5m × 0.75m (length × width) was mounted onto one corner of the chamber on which the testing device was placed, and the test performed. A transparent plexiglass of measurement 54 x 80 cm (width x length) was placed dividing the table into two sides: side A and side B **(Figure 1.A)**. The demonstrator was seated on a perch adjacent to the testing device on Side A. Only one device (either the ‘door’ or the ‘plug’) was available to the birds at a time. On Side B, the test bird was placed on a perch to observe the demonstrator. The experimenter stood on the side of the demonstrator (side A) to place the reward in the reward box, in clear sight of the demonstrator and the subject, followed by the demonstration. After the demonstration, the experimenter switched the sides of the demonstrator and the test subject to begin the test trial. The baseline control setup and procedure were identical to the test setup, except that the control subjects were tested without a demonstrator present.

**Figure 1:**
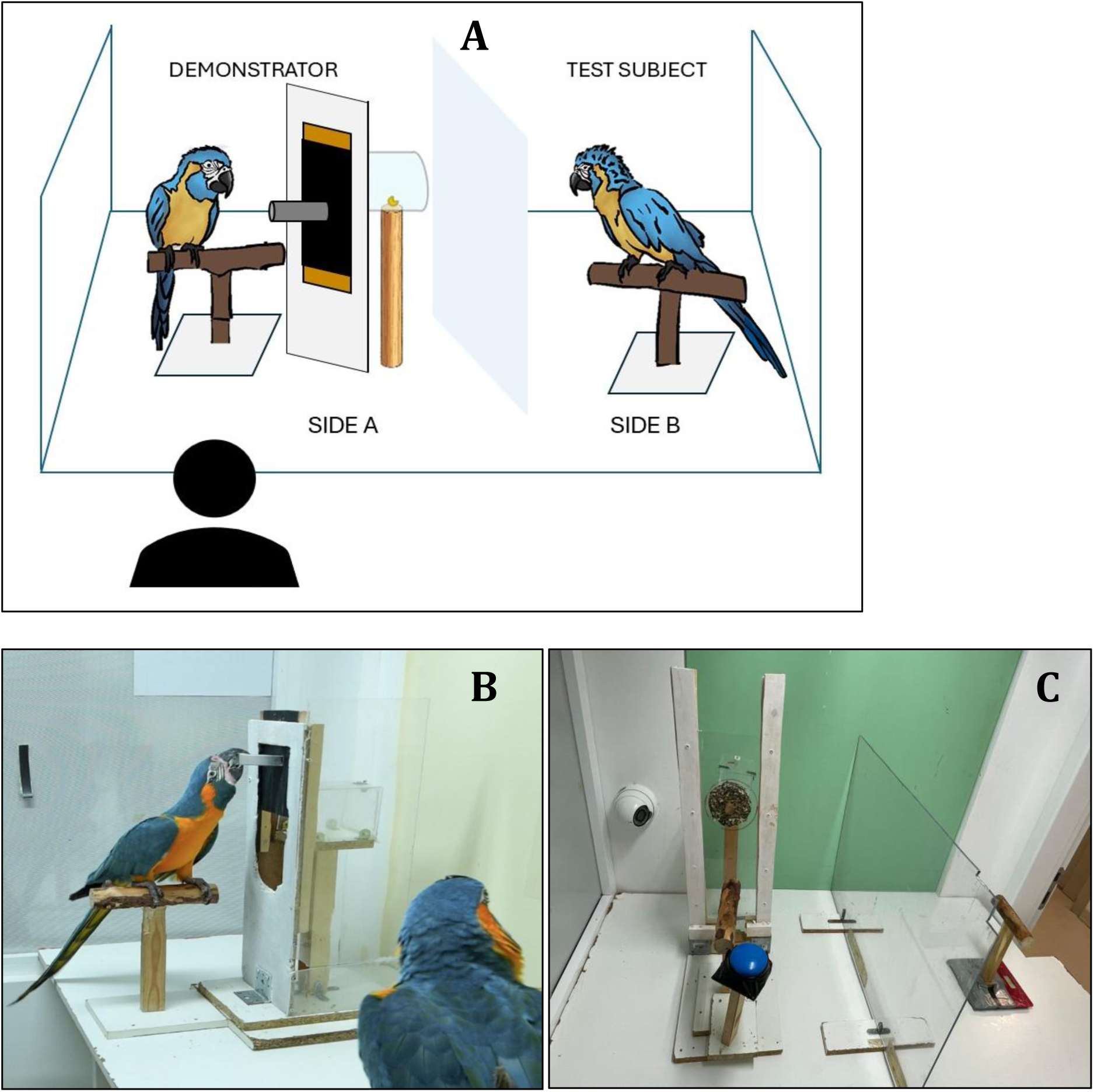
A. Representation of the experimental setup. The test device is on side A, and the test subject’s perch is on side B. The experimenter stands beside the demonstrator/model on side A. The two sides are divided by a plexiglass screen. **B.** View of the ‘*door*’ device with the demonstrator performing *UP* and the test bird observing from side B. **C.** View of the ‘*plug*’ device in the experimental chamber.

### Test apparatus

#### 1. ‘Door’ device

The ‘door’ device was developed from the social learning study conducted on chimpanzees by Watson and colleagues (Watson et al., 2018). The problem-solving task involved pushing UP *or DOWN* a sliding panel (17 x 8cm), mounted against a rectangular window of 23 x 8cm carved on a vertical frame of dimensions 55 x 14 cm (height x breadth) at a height of 23cm from below (see **Figure 1.B)**, hence presenting a bi-directionally controlled task. The sliding panel was fitted as an opaque door within the window, leaving a gap of 3cm above and below the panel. A knob (8cm) was attached to the centre of the sliding panel for easy manipulation. The panel could be moved up or down, either by grasping the knob or by accessing the 3cm gaps above and below the panel. A stable perch of 25 cm height was placed close to the device, on which the parrot could sit and easily access and operate the sliding panel. A transparent reward box was positioned behind the door, at a gap of 3 cm, from which the subject could obtain the reward once it had moved the panel up or down. The reward box could be baited through the 3 cm gap behind the frame, or through the already opened ‘door’ (after the bird had manipulated it). The ‘door’ was constructed to make upward and downward pushing of the panel equally challenging.

#### 2. ‘Plug’ device

The second problem-solving task, a classical two-action task, involved *PULL*ing or *PUSH*ing*’* a cork plug (**Figure 1.C**). A circular hole (5cm diameter) was cut out of a vertical slab of plexiglass (60 x 23 cm, length x breadth), which was supported by wooden frames on each side, at 40 cm height. The hole was plugged with a cork which could be pulled out grasping a knob (8cm) attached to the plug or pushed in with the beak by touching the circular surface area of the cork, in order to make the reward placed in a transparent box 3 cm behind the frame, accessible, similar to the previous device. The reward box could be baited through the 3 cm gap behind the ‘plug’ or through the already exposed hole (after the bird had manipulated the device). A perch of height 25 cm was placed close to the glass slab from which the seated bird could easily access and operate the cork. A blue buzzer was attached to the corner of the perch representing the irrelevant overimitation task.

### General procedure

#### Habituation phase

All 15 birds were habituated to the testing room and the experimental setup for two-three days. The subjects were placed on side B and provided with seeds while the setup was in full view of the subject. Next, they were moved on a handheld perch to the standing perch on side A, adjacent to the experimental devices, where they were fed seeds to habituate them to being near the novel experimental devices. Once they appeared completely relaxed, subjects were trained to retrieve nuts from the open transparent reward boxes of both experimental devices (without any manipulation of the devices). After the subjects readily fed from the open boxes without any hesitation, the habituation was terminated.

#### Control Sessions

Control subjects (N=5) were tested to establish the baseline preference for the species. Each subject was taken into the test chamber and placed on a perch on side B (**Figure 1.A**) as the first trial commenced. The reward was placed in the reward box through the gap at the back of the wooden frame, while the sliding panel on the ‘door’ or the cork on the ‘plug’ device was in their starting (closed) position, while the bird was observing. Then, the naive control bird was moved on a handheld perch and placed on the perch on side A, thus getting access to one of the two experimental devices. Depending on whether there was manipulation (in any direction) or no manipulation within two minutes from the moment the subject had been placed on the perch on side A, the trial was considered successful or failed, respectively. After two minutes, the bird was placed back on side B and the experimenter rebaited the reward box (in case of manipulation) and repositioned the ‘door’ or the ‘plug’ to its starting position, in clear view of the subject. The subject was moved again onto the perch on side A and the 2nd trial began. If a subject did not respond, i.e., attempt to touch or manipulate any part of the device in the first two trials, the experimenter switched to the second device for the following test trial to avoid frustration with this particular device. Each session ended after a maximum of three trials with each device or after the first two consecutive failed trials for each device. A total of 8 sessions were conducted. Each session consisted of up to six trials in total (up to 3 for each device) if successful.

#### Demonstrator Training

Two birds, Lady and Sherlock (from the control group, see **Table 1**) were chosen as demonstrators. They were trained to perform first the irrelevant task and then the relevant task in a sequence for each experimental device. Training relied on gradual shaping with positive reinforcement and operant conditioning. Both Lady and Sherlock were trained to perform the non-preferred direction/action in the relevant two-action tasks. For the ‘door’ device, the irrelevant task involved rotating on the perch in response to a specific hand command from the experimenter (see Supplementary Information). For the ‘plug’ device, the irrelevant task required pressing a buzzer with the foot only, producing a sound. After the demonstrators were given a verbal cue (“GO”) to perform the relevant two-action task. Training continued for up to two weeks, until both birds reliably performed the sequence of tasks.

#### Test sessions

N=10 subjects (5 adults and 5 subadults) were tested in the presence of one of the two demonstrators. Each trial consisted of two demonstrations followed by a test. The demonstrator was first placed on side A and the test bird on side B for observing the demonstration twice (Figure 1.A). The reward was placed in the reward box through the 3cm gap at the back of the frame or through the already opened ‘door’ or ‘plug’, in clear view of the demonstrator and the subject. The demonstrator was signalled by the experimenter to perform the trained sequential task for the respective device and take the reward. Following the second demonstration, the positions of the test subject and the demonstrator were interchanged using handheld perches. While the device was being baited through the already opened slot (from the previous demonstration), the test subject was seated on a perch, at a slight distance from where it could not access the device. When the panel and the cork on the ‘door’ or ‘plug’ was repositioned to the starting position, the perch of the test bird was moved adjacent to the device for the test trial to commence. The subject was allowed to manipulate the setup for a maximum of two minutes (120 seconds), after which the test trial concluded, and the next trial began with another demonstration following the same steps. Each session consisted of a maximum of three trials per device. If the subject failed to interact with one of the two devices in the first two consecutive trials, the device was alternated with the other, or, if it happened with the second device, the session was terminated to avoid frustrating the bird. We conducted a total of eight test sessions with a maximum of 24 trials for each device.

#### Video analysis and data scoring

The video coding for the tests and controls was performed using Solomon coder version 19.08.02 (©2015 by András Péter). The actions performed-*UP*, *DOWN*, *PUSH* or *PULL* – each was coded as a successful trial (trial in which the device was opened, and the reward was obtained) and failure of manipulation of the device was coded as an unsuccessful trial. The success rate for *UP, DOWN, PUSH* and *PULL* were separately calculated for each subject as the total success score divided by the number of trials. If a subject’s success rate for *UP* vs. *DOWN* or *PUSH* vs. *PULL* exceeded 50%, that direction/action was assigned as its action preference; if the rates for *UP* and *DOWN* or *PUSH* and PULL were approximately equal to 50%, the preference was classified as mixed. For comparing copying vs. non-copying, successful manipulation was separately scored as 1 for matching the demonstrator’s respective action and 0 for failing to match the action. The matched response rate was calculated as the sum of the matched responses divided by the total number of trials for each bird, to determine the overall success scores for copying. The number of trials taken to first successfully manipulate any device was noted as a measure of neophobic tendency. Two different latencies were recorded: i) the time duration from ‘start’ of a trial to the ‘first touch’ of the knob (‘s-tk’) attached to the ‘door’ or the cork ‘plug’ and ii) the time duration from ‘first touch’ to ‘end of manipulation’ (‘tk-man’) when the bird had opened the device to access the reward in the box. Two types of latencies were considered to include all trials, where no manipulation occurred, but the birds approached and touched the device. The mean latencies were calculated as the sum of latencies divided by the total number of trials for each subject. Additionally, relevant overimitation behaviours, i.e., rotating on the perch and pressing the buzzer with the foot were noted during each trial.

#### Statistical analysis

All Statistical analyses were performed using R version 4.2.2 (R Core Team (2021). A one-sided Fisher’s test was performed to investigate whether there was a significant difference between the number of control subjects and test subjects manipulating the ‘door’ *UP* or *DOWN* and for the ‘plug’, *PUSH* in or *PULL* out. A Fisher’s test was performed to examine whether there were significant differences between the response of subadult and adult test subjects. To compare copying and non-copying performance, we fitted generalized linear mixed models (GLMM) with binomial error distribution and logit link function in the *lme4* package of R via *glmer* function, to predict ‘successful manipulation’ in the ‘door’ and the ‘plug’ device. We examined the fixed effects of Group (Test vs. Control) and Age Group (Adult vs. Subadult) on the likelihood of producing a matched response to the demonstrations (scored as 1) vs. baseline response (scored as 0), which is the same as the control group, while including individual identity (Subjects) as a random intercept. All model assumptions (i.e. normality of residuals, presence of outliers, dispersion) were checked using the DHARMa package in R, in addition to multicollinearity. The best fit models were selected on the basis of least AIC scores from the Akaike Information Criterion. To test whether the age of the subjects had any effect on the two types of mean latencies in the test and control group, we fitted generalized linear models (GLMs) with age, group and age-group interaction as predictors for the mean latency to touch and the mean latency to manipulate for each device. We fitted generalised linear models (GLMs) to predict the two types of mean latencies (‘s-tk’ and ‘tk-man’) for the ‘door’ and ‘plug’ devices, with age, groups, and age-group interaction as predictors, as part of exploratory analysis. We excluded the age-group interaction from the models through backward selection and fitted only age and group as the independent variables predicting the mean latencies.

## RESULTS

### Control group

All 5 birds in the control group showed a clear preference, i.e., they uniformly moved the sliding panel of the ‘door’ device *DOWN* and *PULLed* the cork out of the ‘plug’ device. The mean success rate of the five individuals of the group for *UP* was 0.00 and for *DOWN* was 0.73 +-0.11 S.E in the ‘door’ device. The subjects used the gap above the door to move it down, instead of grasping the knob with their beak. The mean success rate for *PUSH* was 0.00 and for PULL was 0.72+-0.04 S.E in the ‘plug’ device. Two control birds used their feet rather than their beak for operating the ‘door’ and the ‘plug’. Before starting to manipulate the device, some control birds attempted to attain the reward by other means such as climbing behind the setup or approaching it from the side without touching the ‘door’ or the ‘plug’. Such exploratory behaviour ceased after the subjects had successfully retrieved the reward from the device for the first time.

### Test group

After establishing the preferred action for each device in the control group, which we took as a proxy for the natural preference exhibited by the study species, two individuals from the control group were trained to demonstrate the opposite, i.e., non-preferred action, which was *UP* for the ‘door’ device and *PUSH* for the ‘plug’ device to the test group. The mean success rate of N=10 individuals of the test group for *UP* was 0.71+-0.12 S.E and for *DOWN* was 0.16 +-0.11 S.E in the ‘door’ device. Significantly more test birds pulled UP (>50% of trials), operating the ‘door’ by grasping the knob with their beak, thus matching the directionality of the movement, as well as the relevant stimulus on the device.

In the ‘plug’ setup, the mean success rate for PUSH was 0.38 +-0.1 S.E and for PULL was 0.52+-0.11 S.E across 10 test subjects. Three of the 10 test birds *PUSHed* the cork in the majority of trials (>50% of trials), thus matching the response of the demonstrators. Five out of 10 test birds consistently *PULLed* the cork like the control group. The remaining two test birds showed both *PULL* and PUSH actions in approximately equal ratio (50% of trials) to retrieve the food from the device and their response was considered as ‘Mixed’ **Table 1** summarizes the performance of the test subjects in the two devices. The test subjects avoided exploratory behaviour of obtaining the reward through other means, which was observed in the control birds.

### Group and age differences in task performance

#### ‘Door’ device

In the ‘door’ device, significantly more test subjects performed the UP action, which matched with the demonstrations, compared to the control group (Fisher’s exact test, p*= 0.0069, 95% CI [0.0, 0.592]), which only pulled *DOWN*. The fitted Generalised Linear Mixed Model (GLMM) was highly significant for group Test (β = 7423.4± 9.45, z = 785.85, p*\** <0.001), correctly predicting the response *UP* that matched with the demonstrations. There was no significant difference between the adults and the subadults (β = 0.12± 8.63, z = 0.01, p=0.98) in their copying performance of the demonstrated action *UP.* **Figure 2.A** depicts the success rate of the test and control groups for the action *UP and DOWN* on the ‘door’ device.

**Figure 2.**
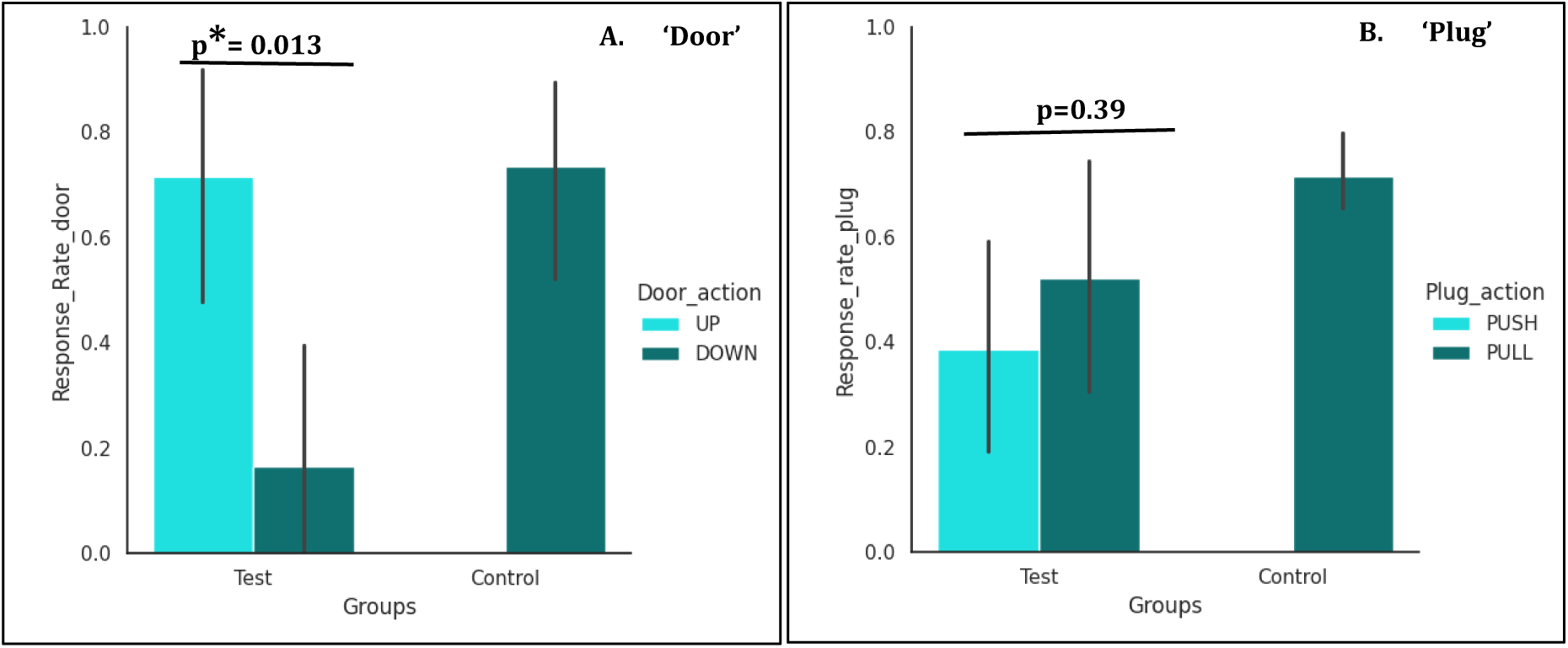
**A: Success rate of test and control subjects for the ‘door’ device.** Light blue represents the response *UP* (matching the demonstrator’s action) and dark green represents *DOWN* (not matching the demonstrator’s action). Only the test subjects performed the action *UP* (the demonstrated action). The success rate of the test birds with the ‘door’ device was significantly higher for the action *UP* (the demonstrated action) compared to the action *DOWN* (Mann-Whitney U test; W=81, p= 0.013). **B: Success rate of test and control subjects for the ‘plug’ device.** Light blue represents *PUSH* in (matching) and dark green represents *PULL* out (non-matching). Only the test subjects performed the action *PUSH* (the demonstrated action). No significant difference was found for the action *PULL* between the test and control subjects (Welch T test; t=1.62, df=11.17, p=0.13). Within the test group, there was no significant difference in the success rate of the action *PUSH* and *PULL* (Welch T test; t= −0.86; df=17.86, p=0.39).

#### ‘Plug’ device

In the ‘plug’ device, there was no significant difference between the number of test subjects performing *PUSH* vs. *PULL* of the plug (Fisher’s exact test, p=0.2, 95% CI [0.0, 0.592]) compared to the control subjects, who only *PULL*ed consistently. The fitted GLMM to predict the matching response to the demonstrations *PUSH* was non-significant for Test Group (β= 20.95 ± 1024.42 SE, z= 0.02, *p* = 0.98). However, there was a significant effect of Age Group: subadults (β= –2.34 ± 1.17 SE, *z* = –2.00, p*\** = 0.045) in predicting *PUSH* responses, indicating that subadult macaws are less likely to match the demonstrations (*PUSH*) in the two-action task than adult macaws, who are more likely to match the demonstrated action. The success rate of the subjects in test and control groups for *PUSH* and *PULL* are given in **Figure 2.B.**

The number of trials taken by the subjects to successfully manipulate each device for the first time and gain rewards did not differ significantly between the test and the control group (see Supplementary Information). The performance of the adults and the subadults in both groups also did not differ significantly in terms of the number of trials taken to first manipulate the devices (Supplementary material).

### Group and age effect on mean latencies

#### ‘Door’ device

**Figures 3.A** and **3.B** illustrate the effect of age and group on the response latencies for the ‘door’ device. For the latency (‘s-tk’) from ‘start’ to ‘touch knob’, the generalised linear model was close to significance (R^2^=0.28, F_2,12_=3.7, p=0.05), with age predicting the mean response time required to approach and touch the setup (β =2.4, p*=0.03) but not groups. A similar result was obtained for the latency (‘tk-man’), from ‘touch knob’ to ‘manipulation’, with an overall significant model (R^2^=0.4, F_2,12_=0.67, p*=0.01) and age predicting the mean latency to manipulate (β =3.0, p*=0.01). Thus, sub-adult birds of both groups took less time to approach and manipulate the ‘door’ device than the adult subjects. Both latencies decreased with an increase in trial numbers in the test and the control group (see Supplementary Information).

**Figure 3:**
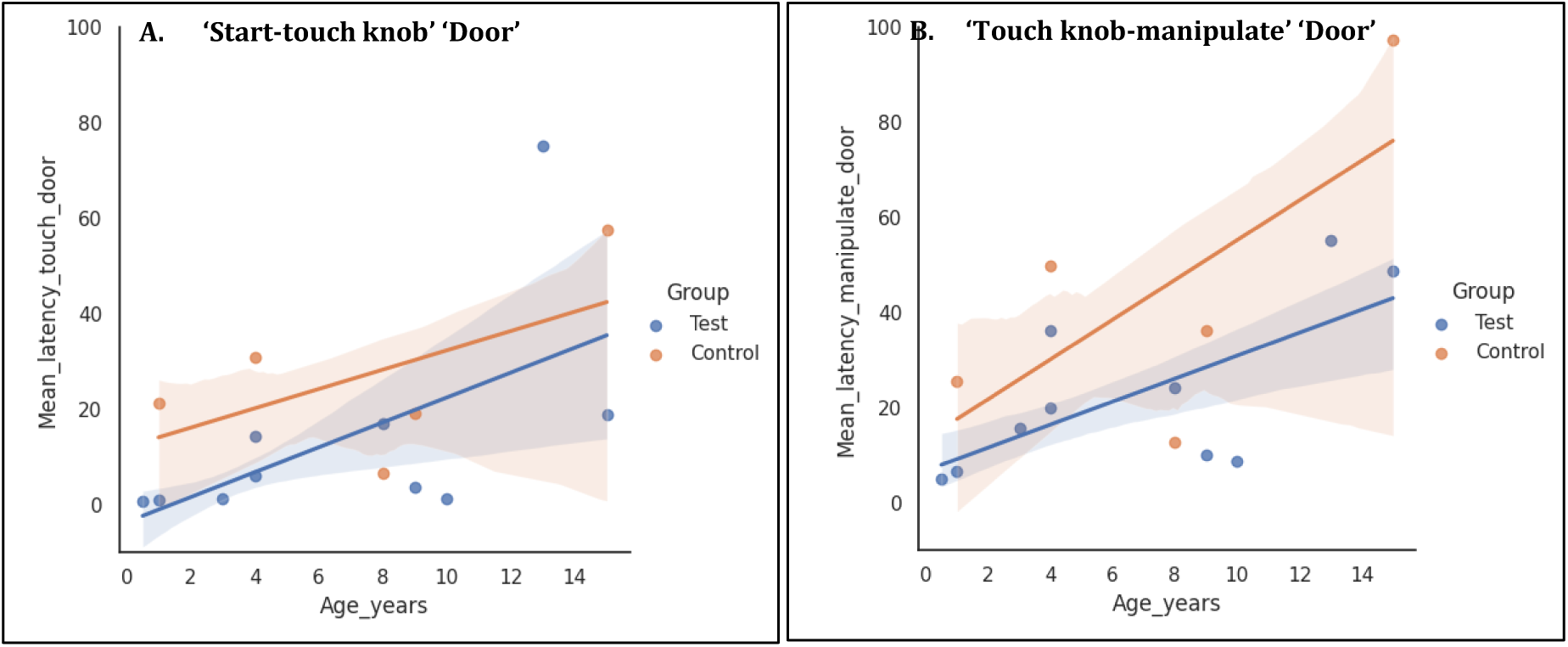
**A:** Regression plot of age (x axis) and group predicting mean latency to approach the ‘door’ device (y axis – duration from start to touch knob) GLM: p=0.05. C.I 75%. **B:** Regression plot of age (x axis) and group predicting the mean latency from touch knob to manipulation (y-axis-duration from touch knob to manipulation end). GLM: p*=0.01. C.I 75%

#### ‘Plug’ device

The generalised linear model with age and group as predictors was overall significant (R^2^=0.3, F_2,12_=5.2, p*=0.02) with group (Group test: β = −19.4, p*=0.03), but not age predicting the mean latency (‘s-tk’) from ‘start’ to ‘touch-knob’ in the ‘plug’ device (**Figure 4.A)**. Thus, the test group took significantly less time to approach and touch the apparatus than the control group, irrespective of the birds’ age. For the second latency (‘tk-man’), ‘touch-knob’ to ‘manipulate’, the fitted regression model was highly significant (R^2^=0.4, F_2,12_=7.2, p*=0.008), with both group (Group test: β = −22.4, p*=0.01) and age (β = 2.05, p*=0.02) predicting the mean time required to successfully manipulate the apparatus, as illustrated in **Figure 4.B**. Thus, in the ‘plug’ device, the control subjects as well as the adult birds took significantly longer time to manipulate the two-action task on the ‘plug’ device.

**Figure 4.**
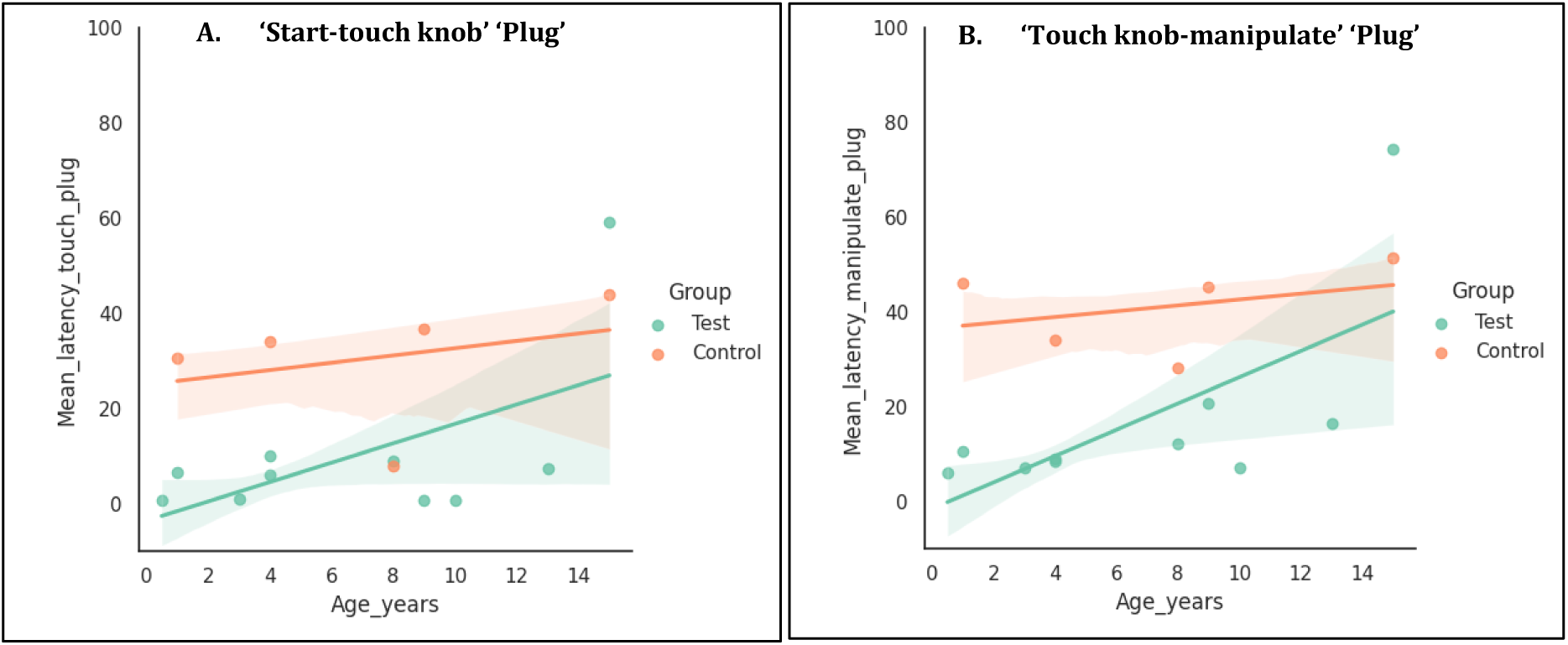
**A:** Regression plot of age (x axis) and group predicting mean latency to approach the ‘plug’ device (y axis – duration from start to touch knob). GLM: p*=0.02, C.I 75%. **B:** Regression plot of age (x axis) and group predicting mean latency from touch knob to manipulation (y-axis-duration from touch knob to manipulation end). GLM: p*=0.008, C.I 75%.

### Performance in overimitation tasks

Out of 10 test birds, three exhibited one of the irrelevant tasks in the first three trials of the experiment. Two of the adult test birds (Wanda and Gargamel) performed a spin, as demonstrated with the ‘door’ device, but did not manipulate or perform the relevant task in the consecutive steps of the trials. One of the subadult test birds (Carrot) pressed the buzzer with its foot in three trials, in the sequence as demonstrated, before manipulating the knob by ‘*PULLing’* (the non-demonstrated action). Thus, it imitated the irrelevant task but did not imitate the relevant task. All subjects had probed the buzzer with their beaks in initial trials, in both test and control conditions, as part of exploratory behaviour.

## DISCUSSION

Our experiment shows that blue-throated macaws can learn to solve physical problem-solving tasks by *copying* their conspecifics. This is implied by the significant effects of conspecific observation on matching the directionality in the bi-directional task (‘door’ device) and the demonstrated action in the two-action task (‘plug’ device). In both tasks, either the majority (in the bi-directional task) or half of the subjects (in the two-action task), showed the directionality/action the demonstrator had shown, which differed from the naturally displayed baseline response exhibited by the control group.

When tested with the bidirectional ‘door’ device, the majority of test birds (eight out of ten) that had watched a demonstrator pulling the door *UP* with their beaks also pulled the door *UP* to retrieve the reward consistently throughout all the sessions. They also used the same technique of grasping the knob with their beaks to manipulate the sliding panel like the demonstrators. In contrast, all the control group birds that had been tested without any demonstrations showed a clear preference for pushing the door *DOWN* with either beak or foot, reflecting their natural response bias towards the device. Our finding that the test group matched the demonstrator in the directionality of solving the bi-directional ‘door’ device does not allow us to infer that *imitation* is the underlying social learning mechanism. *Object movement reenactment*, an emulative process (Custance et al., 1999; Whiten & Ham, 1992), where the subjects learn about the direction of the movement of the object, i.e. in our case the upward movement of the door, is an alternative explanation, which we cannot exclude. Due to our limited sample size, a ghost control group, which would see the *UP* manipulation via an invisible mechanism, in the absence of the conspecific model, as implemented e.g. by Fawcett and colleagues (2002) could not be included given our limited sample size. Failure to copy the demonstrated action in such a ghost control condition would have clearly proven imitation as the underlying mechanism of social learning. Therefore, it was not possible to distinguish between *imitation* and *object movement reenactment* from the bi-directional task. Thus, from the bidirectional task, we conclude that blue-throated macaws *copy* conspecifics through either *object movement reenactment*, an emulative process, or *imitation* of object-directed, transitive actions in the problem-solving task.

When tested with the ‘plug’ device, an actual two-action task, the control group never showed the *PUSH* action, clearly implying *PULL* as a natural tendency of the birds. Five out of 10 test subjects ‘*PUSHed’* in the cork in the same manner as their demonstrator either consistently throughout the test trials or approximately in half of the trials, which none of the subjects in the control group did. This suggests that some subjects indeed learned to solve the ‘plug’ device through *imitation of* the demonstrated non-preferred action in the two-action task. Voekl and Huber (2000), argued in their two-action task experiment with marmosets, that social learning can be considered *imitation* if subjects in the test group perform an action that deviates from their natural tendencies and is observed only in this observation group. In any case, the response of the test birds in the two-action ‘plug’ device less clear than their response in the bidirectional ‘pull’ device. Galef et al. (1986) had revealed that budgerigars used the demonstrated technique: push, to remove a cover from a feeding cup for the first two or three times only in a two-action task, and then switched to the non-demonstrated method: pull, a preferred natural tendency for such action, that overrode the socially learnt technique. The reasons behind this outcome could be linked to the strong tendency in parrot species to “pull” or detach the cork and hold it with their beak or foot (Galef et al. 1986), a behaviour supported by our baseline control group, all of whom preferred to pull out the plug. This suggests that, once the task has been socially learned through overcoming neophobia, the birds prefer to use the most effective method to reach the goal successively.

When examining the performance of the adults and the subadults in the two problem-solving tasks, no significant difference was found in matching the demonstrated action *UP* in the bi-directional ‘door’ device between the two age groups. A meta-analysis of life-history traits affecting social learning in experimental foraging contexts in vertebrates (covering several primate and bird species) showed that age at first reproduction had no discernible effect on social learning (any mechanism) (Penndorf & Aplin, 2020) of the task, implying that social learning of foraging skills is not limited to a narrow age window for most animals (unlike black rats, (Aisner & Terkel, 1992; Zohar & Terkel, 1995)). Most species included in the meta-analysis were large-brained, long-lived taxa such as primates and corvids, which rely on social learning across their lifespan. Parrots exhibit similar life history traits to corvids and primates (Lambert et al., 2019), and the results of our current study corroborate the findings by revealing no difference in the likelihood of matching the demonstrated directionality in the bidirectional ‘door’ device between subadult and adult macaws, indicating that *copying* (or *emulation*) of foraging techniques may be equally important for subadults and adults alike.

However, when we examined the effect of age on social learning further, the subadult macaws were significantly less likely to *imitate* the demonstrated action *PUSH* in the two-action task (‘plug’ device) than the adult macaws, although their *copying* tendency revealed no difference. Miklósi (1999) pointed out that with cognitive maturation by adulthood, animals are more likely to perform advanced social learning like *imitation,* which requires complex cognitive processing, over simpler enhancement or motivational mechanisms like *social facilitation*. Custance *et al*. (1995) argued that they had used chimpanzees older than 4 years of age in their ‘do-as-I-do’ study, because they were old enough to be cognitively capable of copying the demonstrated gestures.’ We found a similar tendency in macaws when manipulating the ‘plug’ device with the two-action task. The subadults were faster to manipulate the ‘plug’ device in the presence of the conspecific than in the control group, thus clearly motivated by the presence of the adult model (*social facilitation*), but they were less likely to match the demonstrated action of the model in the two-action task. Thus, taken together, our findings suggest that while social learning is prevalent and probably adaptive in all age groups of macaws, high-fidelity social learning, like imitation, is more likely to be exhibited by adults when tackling instrumental tasks.

To examine a possible effect of demonstrations on the neophobic tendency of parrots, we compared the latency to touch and manipulate the devices of the control group, which had not received social demonstrations, with that of the test group, which had received demonstrations. It was notable that the test birds took far less time to approach and manipulate the ‘plug’ device compared to the baseline group. This corroborates the *social facilitation* hypothesis that the mere presence of a conspecific motivates the observer (Zentall, 2004, 2006) to interact with a physical problem. It also supports the view that social learning saves time by reducing both neophobia and exploration through trial-and-error learning, leading to faster problem-solving (Auersperg et al., 2014, 2015). Although a similar tendency was observed with the ‘door’ device, the difference between the test and the control group was not significant. However, the subadults approached and manipulated the devices faster than the adults in both devices, suggesting greater exploratory tendency and less neophobia (avoidance of novel objects) (Greenberg & Mettke-Hofmann, 2001) in subadults, which leads to faster acquisition of the foraging techniques.

Overall, our findings support that macaws are not only capable of imitating intransitive actions (no object involved), as evidenced from earlier studies in macaws (Haldar et al., 2024, 2025), but also *copy* and possibly also *imitate* transitive instrumental tasks. While imitation of intransitive actions may serve social functions by fostering group cohesion (Haldar et al., 2024; Lakin & Chartrand, 2003), copying transitive actions is likely to confer adaptive advantages by enabling the transfer of foraging techniques from knowledgeable to naïve individuals. Social learning has been hypothesised as a key phenotypic trait in parrots, for managing the dietary constraints imposed by heavy reliance on unripe fruits that may contain toxins and hard-shelled seeds, which may require extractive foraging (Bradbury & Balsby, 2016). Our findings substantiate this hypothesis by showing that macaws *copy* conspecifics in problem-solving tasks that require learning novel techniques to obtain food. Although little is known about the ecology of blue-throated macaws, they are recognised as specialist foragers on the fruits of Motacú palms in the Bolivian grasslands (Herzog et al., 2021). Like many parrot species, they live in fission– fusion societies that are believed to facilitate the exchange of social information (Bradbury & Balsby, 2016). Copying conspecifics may play a key role in enabling young or inexperienced individuals to acquire “foraging lore” (Bradbury & Balsby, 2016) i.e., local knowledge of effective foraging techniques (e.g., extractive foraging in Goffin’s cockatoos, (Mioduszewska et al., 2022)), or novel foraging resources (e.g., bin opening in sulphur crested cockatoos, (Klump et al., 2021)) that may give rise to foraging cultures in the wild.

Our study also aimed to generate some first findings as to whether macaws may show *overimitation*, i.e., the copying of causally irrelevant actions in addition to the relevant task. *Overimitation* has generally been interpreted as a uniquely human phenomenon, in which children either encode the irrelevant task as necessary for goal acquisition or copy both to satisfy social conventions (Lyons et al., 2011). Overimitation has been observed in companion dogs in rudimentary form (Huber et al., 2018), likely shaped by long history of domestication. As macaws are also sensitive to social cues from their conspecifics, imitating even intransitive, non-goal-directed actions (Haldar et al., 2025), likely serving social bonding and group cohesion, *overimitation* may be expected to occur in this species. We found that one test subject performed the irrelevant action (press buzzer with feet) before performing the relevant action (*PULL* out the knob) sequentially in three trials when tested with the ‘plug’ device. Two more birds performed the irrelevant action: spin, in two trials, but then did not perform the actual relevant action on the ‘door’ device. One may argue that the irrelevant actions could be accidental outcomes, but such incidents were not observed in any of the control trials, implying that the demonstration had an impact on the occurrence of the irrelevant actions. The remaining test subjects did not perform the irrelevant actions and only ‘selectively’ copied the causally relevant tasks (Range et al., 2007); thus, our study provides only a weak indication that *overimitation* may have occurred. Future studies may be more successful by altering the experimental design, because it is possible that our chosen irrelevant actions were unlikely to be copied. For example, concerning the irrelevant action ‘spin’ in the ‘door’ device, the test subjects may have been confused because they were not given the hand command that they were used to associate with ‘spin,’ but had been given to the demonstrator. Nonetheless, to our knowledge, this is the first indication of *overimitation* in an avian species, a behaviour so far tested only in mammals.

The main limitation of this study stems from the small sample size of critically endangered macaws available for the study, which restricted the inclusion of further ghost control and test groups, preventing a clear distinction between imitation and emulative social learning in the bi-directional task. Additionally, the use of gestural commands to execute the irrelevant action by the models may have deterred the overimitation tendency of the test subjects. Future studies should address these points to further corroborate the existence of advanced social learning abilities in macaws.

Overall, our experiment provides evidence that macaws can solve physical problems by *copying* the actions of conspecific demonstrators, thus showing that they are not only capable of imitating intransitive actions (Haldar 2024, 2025) but also can copy transitive actions. We found age to influence social learning in macaws, with different social learning mechanisms being influenced differently by age, confirming that social learning appears important throughout the life span of parrots. Adults seem more likely to imitate conspecifics’ actions, compared to subadults, while subadults appear to respond more readily than adults to simple social learning mechanisms underpinned by motivational or emulative processes. This study also provides first pilot data on overimitation in an avian species, raising the possibility that parrots may exhibit faithful copying of entire action sequences irrespective of their relevance in solving the problem. Our study opens a new line of inquiry into age effects on social learning and overimitation in birds. It encourages future studies to investigate high-fidelity cultural learning using sequences or hierarchically arranged tasks (Whiten et al., 2006) in this important taxonomic group, extending the research beyond primates and mammals to parrots.

## Ethical standards

All applicable international, national, and/or institutional guidelines for the care and use of animals were followed. Under the German Animal Welfare Act of 25th May 1998, Section V, Article 7 and the Spanish Animal Welfare Act 32/2007 of 7th November 2007, Preliminary Title, Article 3, the study was not classified as an animal experiment and did not require any approval from a relevant body. The experiments did not require an application to the Animal Ethics Committee of either Germany or Spain, as animals participated voluntarily in the experiments and were not affected by them in any way. This article does not report any studies with human participants performed by any of the authors. The ARRIVE guidelines for the reporting of animal experiments were followed.

## Acknowledgements

We thank Loro Parque and its president, Mr. Wolfgang Kiessling for their generous support, the access to the birds, and the research facilities. We thank Loro Parque Foundation, its president Mr Christoph Kiessling, and staff, especially the animal caretakers and the veterinary department, for their constant support. We thank Prof. Andrew Whiten of the University of St. Andrews for commenting on the initial drafts of the manuscript. We thank Dr. Sara Torres Ortiz for the images of the parrots. Esha Haldar received funding from DAAD Graduate School Scholarship Program and Animal Minds Project e.V.

## Author contributions

E.H. conceived the study. E.H. and A.B. designed and coordinated the study. E.H. and D.A. collected the data. D.A. coded the videos, E.H and D.A analysed the data. E.H wrote the article. A.B. commented and edited the article. All authors gave final approval for publication and agree to be held accountable for the work performed therein. The authors declare no competing interests.

## Data availability

Raw and analysed data with R codes have been deposited at Figshare and are publicly available at DOI: 10.6084/m9.figshare.30171553

## Materials availability

Detailed description of the materials is listed in the Methods and Supplementary section. Any further information is available from the corresponding authors E.H and A.M.P.B.

## Code availability

This paper reports R codes for known statistical analysis like GLMM, GLM and Fisher’s Exact tests.

