## Supplementary Information for "Subadult macaws (*Ara glaucogularis*) copy better than adults but are less likely to imitate in problem-solving tasks"

**Methods**

**Housing and rearing conditions**

All parrots were group-housed in semi-outdoor aviaries. Measurements of the aviaries were 1.80 × 3.40 × 3 m (width × length × height). All aviaries were equipped with UV-light lamps namely Arcadia Zoo Bars (Arcadia 54W Freshwater Pro and Arcadia 54W D3 Reptile lamp) to ensure sufficient exposure to UV light. Outdoor temperatures during the research period fluctuated between 20 to 26 degrees Celsius during the day and 15 to 21 degrees Celsius during the night with interspersed periods of light raining. The subjects are well habituated to human presence and participate in cognitive testing regularly. Before the experiment commenced the subjects had been habituated to basic handling and clickers, remaining on the perch in the experimental chamber in the presence of a human experimenter and to receiving rewards from human hands.

**Hand commands for irrelevant tasks**

**Fig. S1. Command for irrelevant action 'spin'**

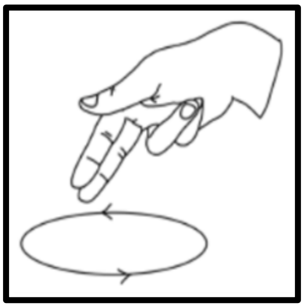

**RESULTS**

**Latencies**

When generalised linear models were fitted to test effect of age as the only predictor for latency of response, the models were significant for both the devices.

**‘Door’ device**

Two types of mean latencies were calculated for each group. For the first latency, the time taken from the ‘start’ of a trial to the ‘first touch’ on the knob of the door (not any other parts of the device) were calculated. The test group took significantly lesser time than the control group ((Mann-Whitney U test: W=42, p*=0.039). There was no difference in the second latency: the time taken from touching the manipulandum till completion of manipulation in the two groups (Mann-Whitney U test: W=37, p=0.16) (**see Figure S2.A).**

**‘Plug’ device**

In the ‘plug’ device (**Figure S2.B**), the time taken from ‘start’ of the trial till first ‘touch’ on the knob, was found to be significantly more in the control group than in the test group watching a demonstrator (Mann-Whitney U test: W=43, p*=0.03). Similar results were obtained for the second type of latency: from ‘touching’ the knob (manipulandum) till end of ‘manipulation’, with control birds taking significantly longer time than the test birds (Mann-Whitney U test: W=4, p*=0.01).

**Figure S2: Effect of group (controls and tests) on latencies of manipulation in the ‘door’ device (A) and the ‘plug’ device (B). Mann-Whitney analysis of average latencies of start to tk (touch knob) and touch knob to manipulation between control and test birds. Pink represents the average latencies of the test birds and red represents the average latencies of the control birds.**

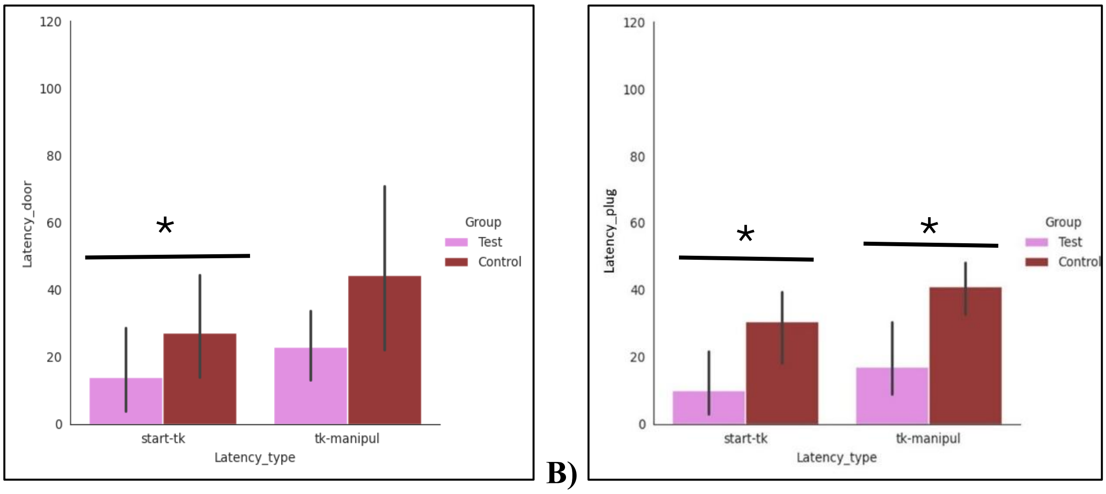


A) B)

**Trial of first manipulation**

The number of trials taken by the subjects to successfully manipulate the devices for the first time and gain rewards did not differ significantly between the test and the control condition. The performance of the adults vs. subadults in the test condition also did not differ significantly. Generalised linear models fitted to test age and group as independent predictors of 'trial of first manipulation', were non-significant for the 'door' device ($R^2$=0.1, $F_{2,12}$=1.8, p=0.2) as well the 'plug' device ($R^2$=0.7, $F_{2,12}$=1.5, p=0.25).

**Latencies across sessions**

We further tested whether there were differences in the time taken for manipulation for test and control birds across the sessions. **Figures S3 and S4** show mean latencies for the time taken to approach the device (start to touch knob) and time taken to complete manipulation (touch knob to manipulation in the control and test birds across all sessions for both devices. The time taken to approach and manipulate decreased across sessions for both controls and test birds. The test birds had lower latencies compared to the controls.

**Figure S3: Illustration of average latencies (y-axis) per session (x-axis) in control and test birds across all sessions in the 'door' device. Pink: control, Green: test birds. A) Average latencies from start to touch knob. B) Average latencies from touch knob to manipulation end**

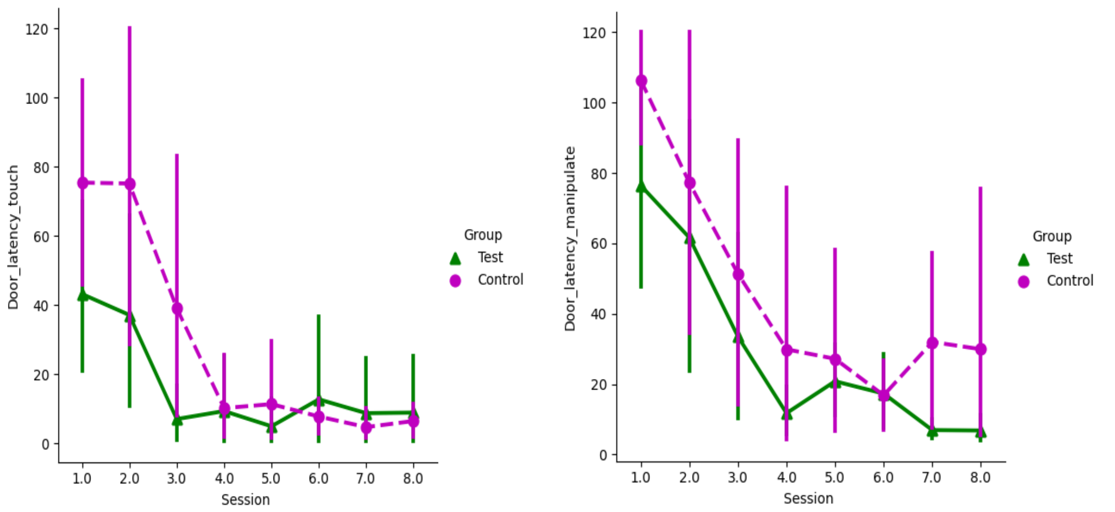


A)

**Figure S4: Illustration of average latencies (y axis) per session (x axis) in control and test birds across 8 sessions in the 'plug' device. Yellow-control; blue-test birds. A) Average latencies from start to touch knob. B) Average latencies from touch knob to manipulation end**

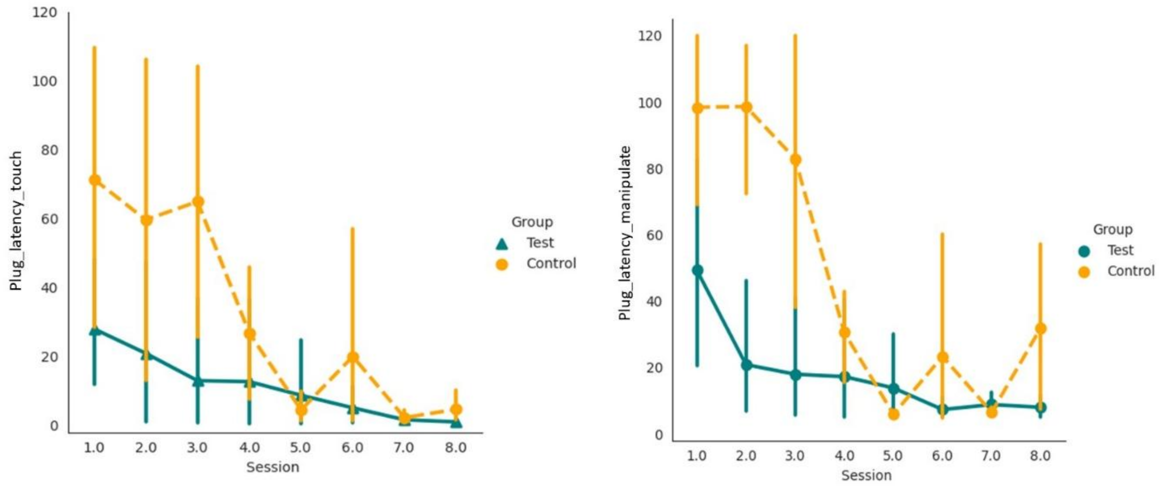


A) B)
